# Fibroblast-derived *HGF* integrates muscle and nerve development during morphogenesis of the mammalian diaphragm

**DOI:** 10.1101/2021.09.29.462475

**Authors:** Elizabeth M. Sefton, Mirialys Gallardo, Claire E. Tobin, Mary P. Colasanto, Gabrielle Kardon

## Abstract

The diaphragm is a domed muscle between the thorax and abdomen essential for breathing in mammals. Diaphragm development requires the coordinated development of muscle, connective tissue, and nerve, which are derived from different embryonic sources. Defects in diaphragm development cause the common and often lethal birth defect, Congenital Diaphragmatic Hernias (CDH). HGF/MET signaling is required for diaphragm muscularization, but the source of HGF and the specific functions of this pathway in muscle progenitors or potentially the phrenic nerve have not been explicitly tested. Using conditional mutagenesis and pharmacological inhibition of MET, we demonstrate that the pleuroperitoneal folds (PPFs), transient embryonic structures that give rise to the connective tissue, are the source of HGF critical for diaphragm muscularization and phrenic nerve primary branching. HGF not only is required for recruitment of muscle progenitors to the diaphragm, but is continuously required for maintenance and motility of the pool of progenitors to enable full muscularization. Thus, the connective tissue fibroblasts and HGF coordinately regulate diaphragm muscularization and innervation. Defects in PPF-derived HGF result in muscleless regions that are susceptible to CDH.

**Summary Statement:** Fibroblast-derived HGF signals to Met+ muscle progenitors and nerve to control the expansion of diaphragm muscle and primary branching of phrenic nerve axons - structures critical for breathing in mammals.

## Introduction

The diaphragm is an essential skeletal muscle and a defining feature of mammals (Perry et al., 2010). Contraction of the diaphragm, lying at the base of the thoracic cavity, powers the inspiration phase of respiration (Campbell et al., 1970). The diaphragm also serves an important passive function as a barrier separating the thoracic from the abdominal cavity (Perry et al., 2010). Respiration by the diaphragm is carried out by the domed costal muscle, composed of a radial array of myofibers surrounded by muscle connective tissue, extending laterally from the ribs and medially to a central tendon, and innervated by the phrenic nerve (Merrell and Kardon, 2013).

Diaphragm development requires coordination of multiple embryonic tissues: 1) somites are well established as the source of the diaphragm’s muscle (Allan and Greer, 1997b; Babiuk et al., 2003; Dietrich et al., 1999), 2) the cervical neural tube gives rise to the phrenic nerve (Allan and Greer, 1997a), and 3) the pleuroperitoneal folds (PPFs), paired mesodermal structures located between the thoracic (pleural) and abdominal (peritoneal) cavities form the muscle connective tissue and central tendon (Merrell et al., 2015). Integration of these three tissues into a functional diaphragm is critical, but how their development is coordinated and integrated is largely unknown.

Defects in the development of the diaphragm cause congenital diaphragmatic hernias (CDH), a common (1 in 3,000 births) and costly ($250 million per year in the US) birth defect (Pober, 2007; Raval et al., 2011; Torfs et al., 1992). In CDH there is a defect in the diaphragm’s muscularization and barrier function. As a result, the liver herniates into the thorax, impeding lung development and resulting in long-term morbidity and up to 50% neonatal mortality (Colvin et al., 2005). Correct innervation of the diaphragm by the phrenic nerve is also essential, as breathing must be functional by birth and fetal breathing movements are thought to be important for normal lung development (Jansen and Chernick, 1991).

Recruitment of muscle progenitors and targeting of phrenic nerve axons to the nascent diaphragm are essential first steps for correct diaphragm development. The receptor tyrosine signaling cascade initiated by the binding of the ligand Hepatocyte Growth Factor (HGF) to its receptor Met is as a promising candidate pathway for regulating these steps of diaphragm development, as HGF/MET signaling has been implicated in multiple aspects of muscle and motor neuron development (reviewed by Birchmeier et al., 2003; Maina and Klein, 1999). HGF binding to MET leads to MET phosphorylation and the activation of multiple downstream pathways, including, JNK, MAPK, PI3K/Akt, and FAK (Organ and Tsao, 2011). HGF/MET signaling is a critical regulator of muscle progenitors migrating from somites (Bladt et al., 1995; Dietrich et al., 1999; Maina et al., 1996). HGF is also critical for innervation. In limb development, HGF acts as a chemoattractant and is required for correct guidance of Met+ motor neuron axons as they extend towards the limb and branch to target muscles (Ebens et al., 1996; Yamamoto et al., 1997). Loss of HGF leads to abnormal branching of nerves in the limb (Ebens et al., 1996) and Met signaling is required for distinct functions in different motor neuron pools, including axon growth in the latissimus dorsi and motor neuron survival in the pectoralis minor (Lamballe et al., 2011). In zebrafish, HGF/Met signaling guides vagus motor axons and affects axon target choice (Isabella et al., 2020). Thus, HGF/Met signaling has complex, tissue-specific roles in regulating the neuromuscular system.

Here we dissect the role of HGF/Met signaling in muscularization and innervation of the diaphragm. *Met* mutations have been associated with CDH (Longoni et al., 2014) and *HGF* is downregulated in in mutants or pharmacological treatments that induce diaphragmatic hernias in rodents (Merrell et al., 2015; Takahashi et al., 2016). Furthermore, *Met* null mice lack all diaphragm musculature (Bladt et al., 1995; Dietrich et al., 1999; Maina et al., 1996). However, which cells are the source of HGF and what steps of diaphragm muscle development HGF/MET signaling regulates is unclear. Also, unknown is whether this signaling pathway is important for phrenic nerve innervation. Using conditional mutagenesis, pharmacological treatments, and an *in vitro* primary cell culture system (Bogenschutz et al., 2020), we demonstrate that the diaphragm’s connective tissue fibroblasts are a critical source of HGF that recruits and maintains muscle progenitors into and throughout the developing diaphragm and controls defasciculation of the phrenic nerve.

## Results

### *HGF* and *Met* are expressed in the developing diaphragm

To begin dissecting the precise role(s) of HGF/MET signaling in diaphragm development, we investigated the expression of *HGF* and *Met* in the early diaphragm. We first examined mouse embryos at E10.5 when muscle progenitors are migrating from cervical somites to the nascent diaphragm and progenitors have already populated the forelimb (Sefton et al., 2018). At this stage, *HGF* is expressed in the mesoderm lateral to the somites (but not in the somites themselves) and in the limb bud mesoderm (Fig. 1A), while *Met* is expressed in the somites and the muscle progenitors in the limb bud (Fig. 1D and Sonnenberg et al., 1993). At E11.5 muscle progenitors have migrated into the pleuroperitoneal folds (PPFs) of the diaphragm (Sefton et al., 2018). During this stage *HGF* is expressed throughout the pyramidal PPFs (arrows, Fig. 1B), while *Met* is expressed in a more restricted region in the PPFs (presumably in muscle progenitors, arrows, Fig. 1E), in limb muscle progenitors (Fig. 1E and Sonnenberg et al., 1993), as well as the phrenic nerve (Fig. 1F). By E12.5, the PPFs have expanded ventrally and dorsally across the surface of the liver (Merrell et al., 2015; Sefton et al., 2018). Strikingly, *HGF* is restricted to the ventral and dorsal leading edges of the PPFs (arrows and asterisks, Fig. 1C). Although Met is no longer detectable by whole mount RNA in situ hybridization, qPCR indicates that it is still expressed at E12.5 at comparable cycle threshold values to *HGF* and *Pax7* (Fig. S1). These expression patterns suggest that HGF expressed in the mesoderm adjacent to the somites, PPF fibroblasts, and limb bud fibroblasts activates MET signaling in the diaphragm, limb muscle progenitors and phrenic nerve.

**Figure 1.**
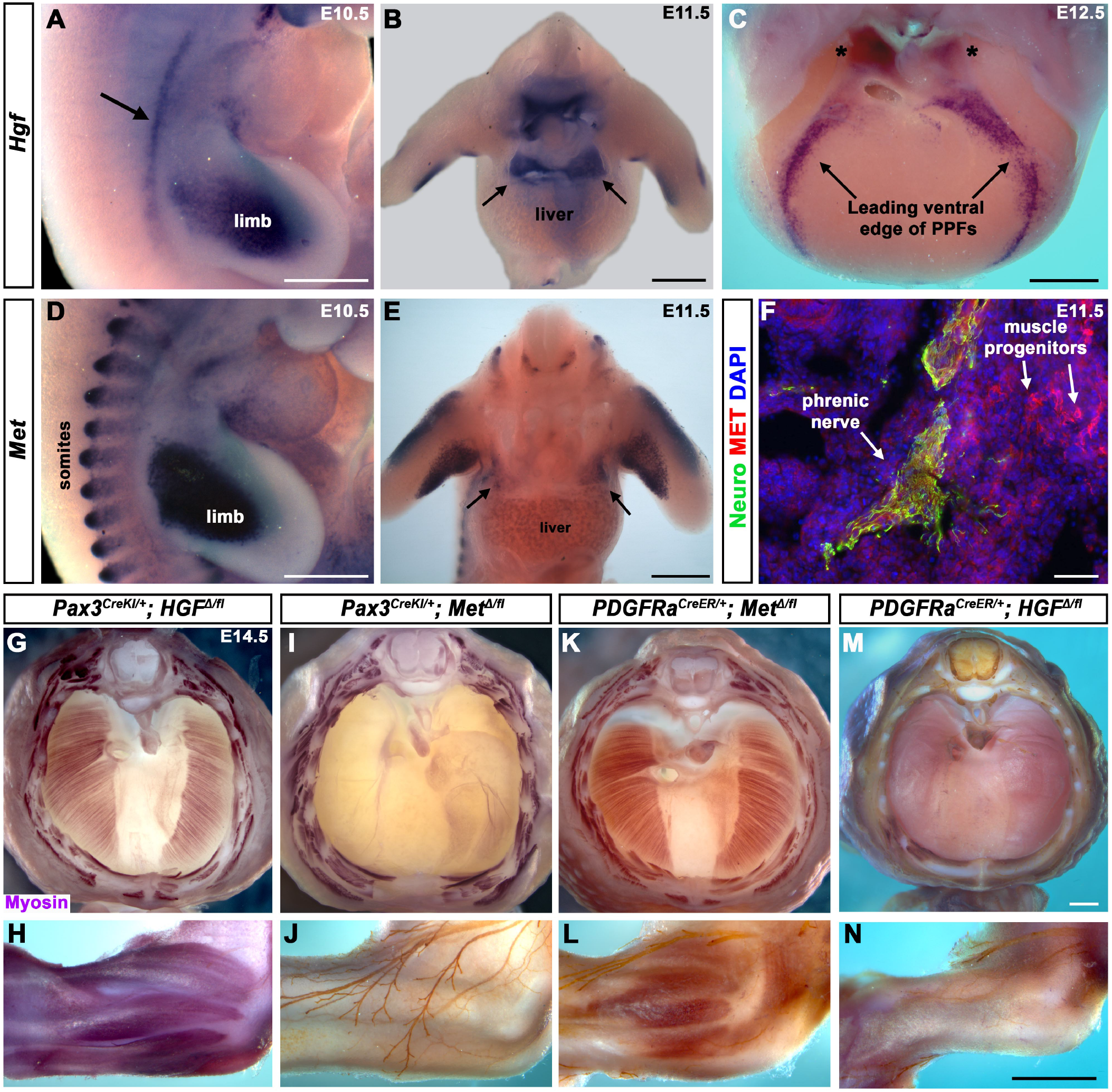
Fibroblast-derived *HGF* and somite-derived *Met* are required for muscularization of the diaphragm and limb. A) Lateral view at E10.5 of *HGF* expression in lateral mesoderm adjacent to somites (arrow) and limb; B) Cranial view of *HGF* expression in E11.5 developing diaphragm and limbs; C) Cranial view of *HGF* expression in E12.5 diaphragm at the leading edges of the PPFs as they spread ventrally (arrows) and dorsally (asterisks). D) Lateral view of E10.5 *Met* expression in muscle progenitors of limb and somites. E) Cranial view of *Met* expression in E11.5 developing diaphragm and limbs. F) Expression of MET and Neurofilament in transverse section through the phrenic nerve at E11.5. G, I, K, M) E14.5 diaphragms stained for Myosin. H, J, L, N) E14.5 forelimbs stained for Myosin and neurofilament. Deletion of *Met* in the Pax3 lineage (I-J) or *HGF* in PDGFRα lineage (tamoxifen at E8.5) (M-N) leads to muscleless diaphragms and muscleless or partially muscularized limbs. Conversely, deletion of *HGF* in Pax3 lineage (G-H) or *Met* in PDGFRα lineage (tamoxifen at E9.5) (K-L) results in normal diaphragm and limb muscle. Scale bars A-E; G, I, K, M: 500 μm; H, J, L, N: 500 μm; F: 50 μm.

### Fibroblast-derived *HGF* and somite-derived *Met* are required for diaphragm and limb skeletal muscle

The complete absence of diaphragm and limb muscles in mice with null-mutations for *Met* (Bladt et al., 1995; Dietrich et al., 1999; Maina et al., 1996) demonstrates that *Met* is critical for development of these muscles. The spatially restricted expression of *Met* and *HGF* suggests that MET signaling in muscle progenitors is induced by HGF in the PPF and limb fibroblasts. Surprisingly, the tissue-specific requirement of Met and HGF has not been genetically tested *in vivo*. We tested whether the receptor is required in somite-derived diaphragm and limb myogenic cells via *Pax3*^*CreKI*^ mice (Engleka et al., 2005), which recombines in the somites, including all trunk myogenic cells, and *Met*^*fl*^ mice (Huh et al., 2004). Consistent with a hypothesized critical role of MET in myogenic cells, conditional deletion of *Met* in the somitic lineage results in a muscleless diaphragm and limbs (Fig. 1I-J). Additionally, we tested whether HGF derived from PPF and limb fibroblasts is critical via *PDGFRα*^*CreERT2*^ mice (Chung et al., 2018). *PDGFRα* is expressed in the PPFs of the diaphragm (Fig. S2 A-D), and *PDGFRα*^*CreERT2*^ drives Cre expression in the connective tissue fibroblasts of the diaphragm and limb, but not in muscle fibers (Fig. S2 E-L). When combined with *HGF*^*fl*^ (Phaneuf et al., 2004), *Pdgfrα*^*CreERT2/+*^; *HGF*^*Δ/fl*^ mice given tamoxifen at E8.5 have a muscleless diaphragm (Fig. 1M) and partial or complete loss of muscle in limbs (Fig. 1N), demonstrating a crucial role for HGF derived from PPF and limb fibroblasts. We also tested the alternative hypotheses that HGF is produced by myogenic cells and Met signaling is active in fibroblasts, but *Pax3*^*CreKI/+*^; *HGF*^*Δ/flox*^ and *Pdgfrα*^*CreERT2/+*^; *Met*^*Δ/fl*^ mice have normal diaphragm and limb musculature (Fig. 1G, H, K, L). In summary, these data establish that HGF derived from PPF and limb fibroblasts induces MET signaling in somite-derived myogenic cells, which is required for muscularization of the diaphragm and the limbs.

### Diaphragm and shoulder muscle progenitors require fibroblast-derived *HGF* during a similar temporal window

HGF/Met signaling is required for both diaphragm and limb muscle, but it is unclear if HGF is required during the same temporal window for development of diaphragm and forelimb muscles. This question is of particular interest because it has been proposed that shoulder muscle progenitors migrate at a similar time as diaphragm progenitors and during evolution recruitment of shoulder muscle progenitors into the nascent diaphragm led to the muscularization of the diaphragm in mammals (Hirasawa and Kuratani, 2013). To dissect the temporal requirement for fibroblast-derived *HGF* for diaphragm and forelimb muscles, we examined these muscles in E16.5-18.5 *PDGFRα*^*CreERT2/+*^; *HGF*^*Δ/flox*^ mice given tamoxifen at E9.5 or E10.5, when muscle progenitors are actively delaminating from the somites and migrating into the nascent diaphragm and forelimb. We initially gave 6 mg of tamoxifen at E9.5, but only two embryos survived. Based on these two embryos, there was no obvious difference between 6mg versus 3mg tamoxifen on muscle: one embryo given 6mg had muscleless limbs and diaphragm (Fig. 2, Q-T), while the other had partial muscle in the diaphragm with normal limb muscle (data not shown). Therefore, we used 3 mg of tamoxifen for all subsequent experiments and compared diaphragm and limb defects in individual embryos (each row in Fig. 2 shows diaphragm and forelimb muscle from a single embryo). In the most mildly affected embryos, the diaphragm is missing a small ventral patch of muscle with normal forelimb muscles (Fig. 2 E-H; n = 5/11). In the most severely affected case, both the forelimb and diaphragm are muscleless (n = 1/11). A small number of mutants had muscleless diaphragms, but normal forelimb muscles (n = 2/11). Strikingly, a subset of mutants had muscleless (or nearly muscleless) diaphragms and displayed specific defects in shoulder musculature (Fig. 2 I-P; n = 4/11). The acromiodeltoid was absent or reduced and mis-patterned (Fig. 2 L, P) and the spinodeltoid was strongly reduced in size, while other forelimb muscles appeared normal (Fig. 2 K, O). When tamoxifen was given at E10.5, diaphragms had partial muscle, with normal forelimb muscles (n = 3/3; data not shown). In summary, muscleless limbs are always accompanied by a muscleless diaphragm, suggesting that these embryos had an early defect whereby muscle progenitors were unable to delaminate from the somites and migrate into nascent forelimbs and diaphragm. Partially muscularized diaphragms are associated with normal limb muscle. While a muscleless or nearly muscleless diaphragm may or may not have accompanying limb defects, loss of shoulder muscle was always associated with a muscleless or nearly muscleless diaphragm. These intermediate phenotypes indicate that most muscle progenitors migrate into the forelimb in advance of progenitors migrating into the diaphragm. However, based on their similar temporal sensitivity to HGF/MET signaling, the shoulder acromiodeltoid and spinodeltoid progenitors migrate at a similar later time as the diaphragm progenitors and thus suggests that the diaphragm shares a closer relationship to shoulder muscles than other forelimb muscles.

**Figure 2.**
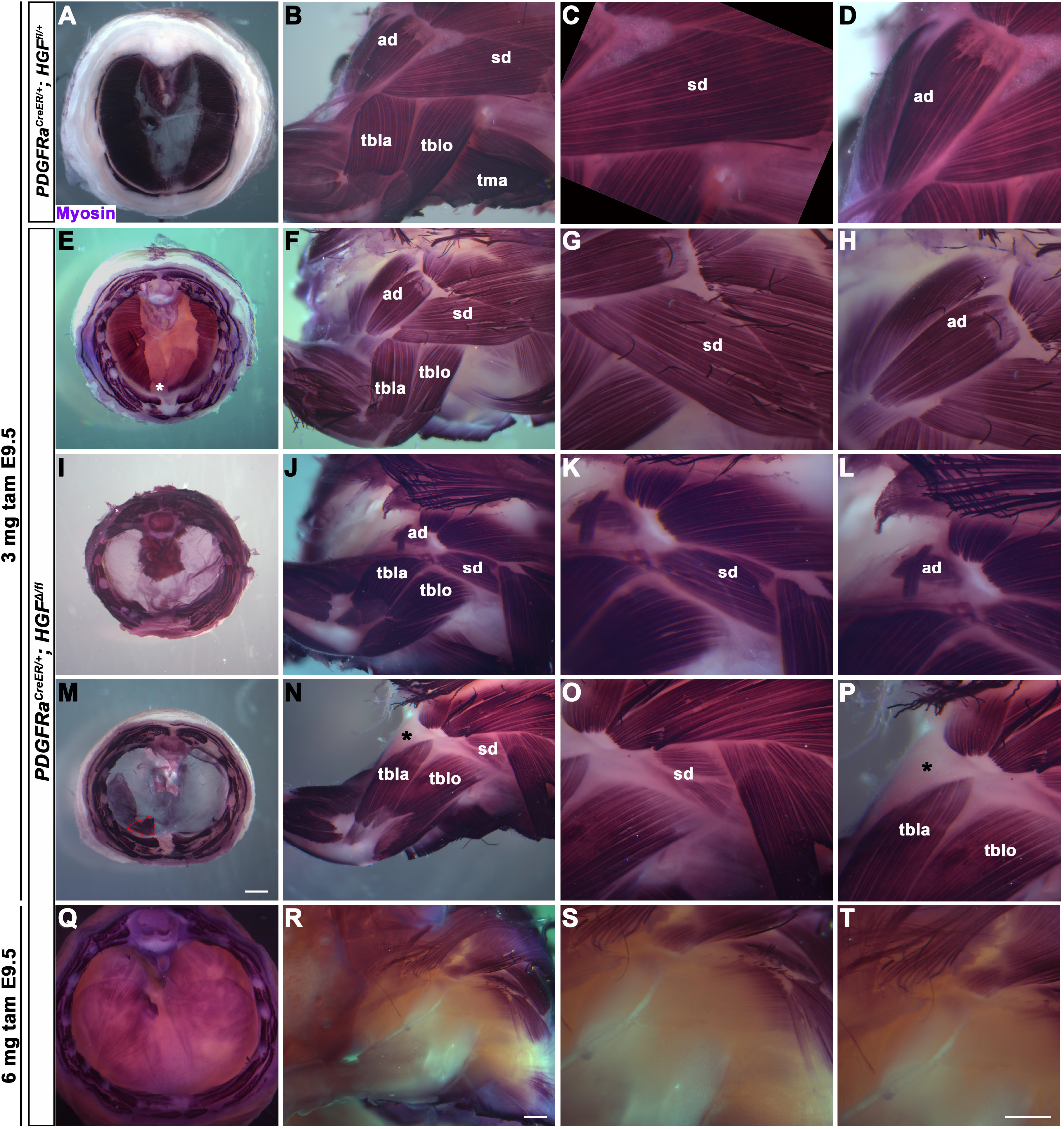
Reduced and mispatterned acromiodeltoid and spinodeltoid accompanies loss of diaphragm muscle following deletion of *HGF* in the *PDGFR* lineage. A-D) Diaphragm and limb musculature in control *PDGFRα*^*CreER/+*^; *HGF*^*fl/+*^ given 3mg tamoxifen at E9.5. E-H, I-L, M-P) Diaphragm and limb muscle in three mutant *PDGFRα*^*CreER/+*^; *HGF*^*Δ/fl*^ embryos given 3mg tamoxifen at E9.5. In the mildest phenotype, loss of ventral diaphragm muscle (asterisk, E) but normal limb and shoulder muscles (F-H; n = 5/11). In the moderate phenotype, shoulder muscles were affected with the diaphragm (n = 4/11). Absence of diaphragm muscle (I) accompanied with normal limb muscle except spinodeltoid reduced and acromiodeltoid reduced and mispatterned (J-L). Similarly, loss of diaphragm muscle, except in ventral-most region (red dotted line, remaining purple is AP stain trapped in connective tissue layer M) and normal limb muscle except spinodeltoid reduced and acromiodeltoid absent (asterisk, N-P). Q-T) In *PDGFR*α^*CreER/+*^; *HGF*^*Δ/fl*^ embryo given 6 mg tamoxifen at E9.5 near complete loss of both diaphragm and limb muscle (n = 1/2). Embryos harvested between E16.5 and E18.5. All samples stained with myosin antibody. ad, acromiodeltoid; sd, spinodeltoid; tbla, triceps brachii lateral; tblo, triceps brachii long Scale bars A, E, I, M: 1mm; B, F, J, N, Q, R: 500 μm; C, D, G, H, K, L, O, P, S, T: 500 μm.

### PPF-derived HGF controls phrenic nerve defasciculation in a dose-dependent manner, independent of muscle

HGF signaling can act as a neurotrophic factor and chemoattractant in spinal motor neurons and cranial axons (Caton et al., 2000; Ebens et al., 1996; Isabella et al., 2020). However, the function of HGF and MET in development of the phrenic nerve, the sole source of motor innervation in the diaphragm, has not been examined. To test the role of HGF/Met signaling in the phrenic nerve, *HGF*^*Δ/Δ*^, *Met*^*Δ/Δ*^, and *Prx1*^*Cre/+*^;*HGF*^*fl/fl*^ mice were stained for neurofilament (Fig. 3). By E12.0, in control mice the phrenic nerve has reached the PPFs and defasciculates into numerous small branchpoints prior to the full extension of the 3 primary branches (Fig. 3A, arrows). However, in *HGF*^*Δ/Δ*^ mutants, while the phrenic nerves reach the surface of the diaphragm, they do not correctly branch and defasciculate. The right phrenic nerve sends out 2 branches in a Y-shape (surrounding the vena cava) instead of numerous small branches (n = 3/3; Fig. 3B) and the left phrenic nerve also fails to defasiculate to the same extent as in control embryos (Fig. 3B). To test if the PPF fibroblasts are a critical source of *HGF* for branching of the phrenic nerve, *HGF* was conditionally deleted using *Prx1Cre*^*Tg*^ (Logan et al., 2002), which robustly recombines in PPF-derived fibroblasts (Merrell et al., 2015). Consistent with *Pdgfrα*^*CreERT2/+*^;*HGF*^*Δ/flox*^ mice, the diaphragms of *Prx1Cre*^*Tg/+*^;*HGF*^*Δ/flox*^ are muscleless (n = 5/12) or partially muscularized (n = 7/12). While the phrenic nerves reach the muscleless diaphragm, they lack primary and secondary branches (arrows, Fig. 3C-D). Thus, PPF-derived HGF is necessary for phrenic nerve defasciculation. Similar defasciculation phenotypes are also present in *Met*^*Δ/Δ*^ mutants. Confocal analysis of E11.5 diaphragms, when the phrenic nerve is just reaching the PPFs, reveals that defasciculation defects are present in *Met*^*Δ/Δ*^ mutants by this early time point (Fig. 3I, J). Comparison of E11.5 *Met*^*+/+*^, *Met* ^Δ/+^, and *Met*^*Δ/Δ*^ diaphragms reveals a dose-dependent requirement for Met, as the number of fascicles is lower in heterozygotes and is further reduced in homozygous mutants (Fig. 3E-K). Importantly, the reduced number of branches in *Met* ^Δ/+^ nerves indicates that the reduced branching is not merely the result of the total loss of muscle, as *Met* ^Δ/+^ embryos have normally muscularized diaphragms (e.g. see Fig. 1K). Altogether these data demonstrate that PPF-derived *HGF* is required for normal defasciculation and primary branching of phrenic nerves, in a Met dose-dependent manner that is independent of the presence of muscle.

**Figure 3.**
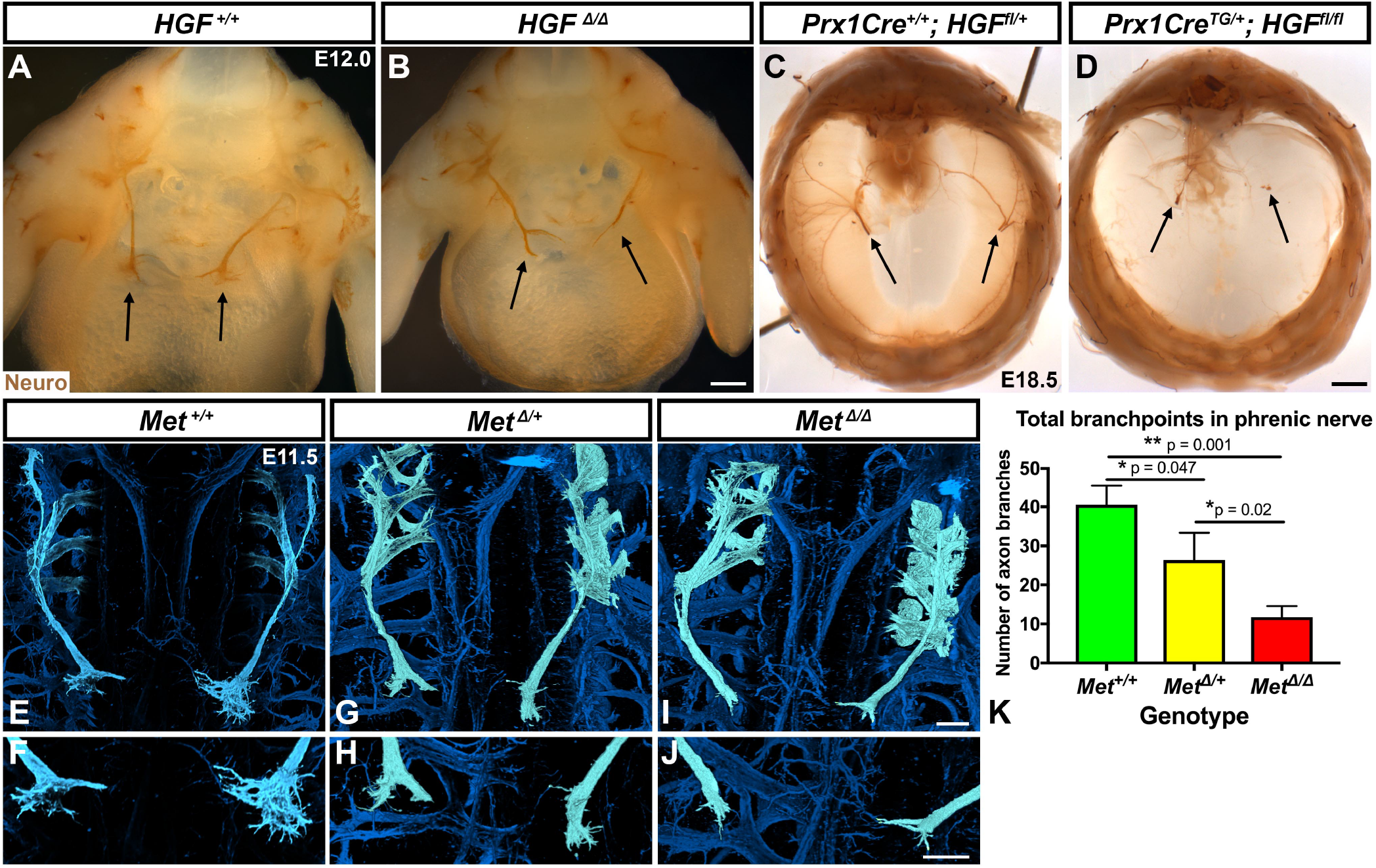
Loss of *HGF* and *Met* leads to defasciculation defects in the phrenic nerve. Wholemount neurofilament staining of the phrenic nerve in control (A, C, E, F) or *HGF* (B, D) or *Met* mutants (G-J). Cranial view of dissected diaphragm region viewed with light microscopy (A-D) showing loss of phrenic nerve branches in *HGF*^/^ diaphragm (arrows, A-B) and *Prx1Cre*^*TG/+*^; *HGF*^*fl/fl*^ diaphragm (arrows, C-D). Dorsal view with confocal microscopy shows reduced phrenic nerve defasciculation in *Met*^*Δ/+*^ and *Met*^*Δ/Δ*^ diaphragms (E-J). Phrenic nerve and C3-5 artificially colored in light blue. D) Quantification of phrenic nerve branchpoints at E11.5 in *Met*^*+/+*^, *Met*^*Δ/+*^, and Met^*Δ/Δ*^ embryos. Scale bars A-B: 250 μm; C-D: 1mm; E, G, I: 100 μm; F, H, J: 100 μm.

### *HGF* is required in fibroblasts to fully muscularized the diaphragm after delamination of muscle progenitors from somites

While HGF/Met signaling is critical for delamination of muscle progenitors from the somites (Dietrich et al., 1999), it is unclear if *HGF* plays a later role in development of the diaphragm’s muscle. To test the later temporal requirement of *HGF* in PPF fibroblasts, we deleted *HGF* via *Pdgfrα*^*CreERT2/+*^; *HGF*^*Δ/fl*^ mice given tamoxifen at different timepoints. When *HGF* via *Pdgfrα*^*CreERT2/+*^; *HGF*^*Δ/fl*^ mice are given tamoxifen at E9.0, prior to the onset of muscle precursor migration to the diaphragm (Sefton et al., 2018), the diaphragm lacks all muscle (Fig. 4B; n = 3/3). This is likely due to a failure of muscle progenitors to delaminate and emigrate from the somites, as in *Met-null* diaphragms (Dietrich et al., 1999). When *HGF* is deleted via tamoxifen at E9.5, when muscle progenitors are delaminating and migrating to the nascent diaphragm, the diaphragm displays large ventral muscleless regions as well as dorsal muscleless patches at E14.5 (Fig. 4C; n = 6/6). Notably, the phrenic nerves in these diaphragms only extends to the regions with muscle (Fig. 4H). When *HGF* is deleted via tamoxifen at E10.5, the muscle reaches its normal ventral extent in most E14.5 embryos (Fig. 4D, n = 11/13). However, when these embryos are allowed to develop to E17.5 (when the muscle has normally expanded to the ventral midline), a large ventral muscleless region persists in mutant embryos (Fig. 4F; n = 3/3). When mutants are given tamoxifen at E11.5, after migration of progenitors to the PPFs has completed, a smaller muscleless region is present in the ventral diaphragm at E17.5 (Fig. 4G; n = 4/4). Thus, these data demonstrate that PPF-derived *HGF* is critical for muscularization of the ventral- and dorsal-most regions of the diaphragm after its initial requirement for muscle precursor delamination from the somites. This role of HGF in the ventral- and dorsal-most regions is consistent with its strong expression in these regions (Fig. 1C).

**Figure 4.**
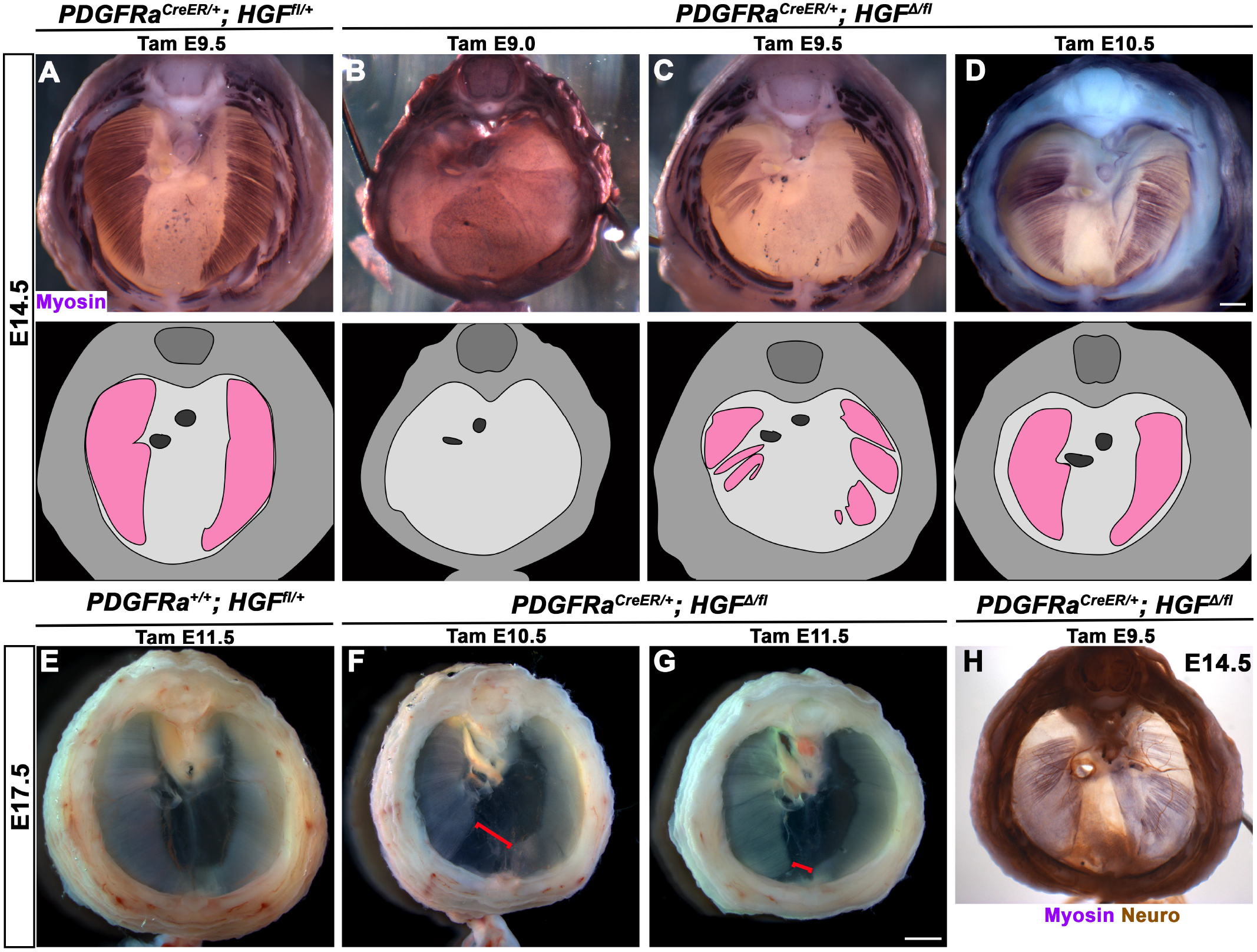
Loss of diaphragm muscle following timed deletion of *Hgf* in the PPF fibroblast lineage. A-D) Upper panels, cranial view of E14.5 diaphragms stained for Myosin. Middle row panels, illustrations of muscle distribution. A) Control *PDGFRα*^*CreER/+*^; *HGF*^*fl/+*^ diaphragm muscle forms two lateral wings that have not yet converged ventrally. B) Muscleless diaphragm when tamoxifen administered at E9.0 (n = 3/3). C) Large muscleless regions in ventral and dorsal diaphragm following tamoxifen administration at E9.5 (n = 6/6). D) Mutant *PDGFRα*^*CreER/+;*^ *HGF*^*Δ/flox*^ muscle typically reaches typical ventral extent when tamoxifen is administered at E10.5 (n = 11/13). E-G) Unstained E17.5 diaphragms in cranial view. E) Control diaphragm muscle has closed ventrally. F) Large ventral muscleless region following *HGF* deletion at E10.5 (red bracket, n = 3/3). G) Smaller ventral muscleless region following tamoxifen at E11.5 (red bracket, n = 4/4). H) Cranial view of E14.5 *PDGFRα*^*CreER/+;*^*HGF*^*Δ/flox*^ mutant with tamoxifen administered at E9.5. Diaphragm stained for Myosin and neurofilament, indicating phrenic nerve tracks with regions of muscle (n = 3/3). Scale bars A-D, H: 500 μm; E-G: 1 mm.

We next sought to determine how PPF-derived HGF regulates development of the dorsal and ventral-most regions of the diaphragm muscle. Our previous studies (Merrell et al., 2015; Sefton et al., 2018) have shown that the PPFs expand dorsally and ventrally, carrying muscle as they expand, and so control overall morphogenesis of the diaphragm. Absence of dorsal and ventral muscle regions in *HGF* mutants could result from a failure of the PPFs to expand dorsally and ventrally and thus lead to the consequent lack of dorsal and ventral diaphragm muscle. To test if the PPF expansion is aberrant following deletion of *HGF*, we examined the diaphragm of *Prx1Cre*^*Tg/+*^; *HGF*^*Δ/fl*^ *Rosa*^*LacZ/+*^ mice, in which we genetically labeled PPFs as they spread across the surface of the liver at E13.5. However, the PPFs reach their normal ventral extent at E13.5 following loss of *HGF* (Fig. S3). This argues that the loss of ventral and dorsal muscle regions is not due to defects in the morphogenesis of the PPFs.

### Development of dorsal and ventral regions of diaphragm muscle requires continuous Met signaling

The experiments conditionally deleting HGF after emigration of myogenic progenitors from the somites indicates that HGF/Met signaling plays additional later roles in development of the diaphragm’s muscle. To specifically test when *Met* signaling is required in myogenic cells, we first deleted *Met* using *Pax7*^*iCre/+*^ or tamoxifen-inducible *Pax7*^*CreERT2*^ mice (Keller et al., 2004; Murphy et al., 2011), which cause Cre-mediated recombination later than *Pax3*^*Cre*^ in a subset of embryonic myogenic progenitors as well as all fetal and adult progenitors (Hutcheson et al., 2009). Neither *Pax7*^*iCre/+*^; *Met* ^Δ/flox^ nor *Pax7*^*CreERT2/+*^; *Met* ^Δ/flox^ embryos displayed any defects in diaphragm muscularization at E14.5 or P1 (Fig. S4). This may indicate that *Met* is not required during fetal myogenesis, as Pax7 is primarily expressed in fetal myogenic progenitors. However, *Met* derived from embryonic Pax3+Pax7-myogenic progenitors is likely present in the muscle of these mutant diaphragms and so does not permit analysis of the consequence of *Met* deletion in muscle.

As an alternate strategy to test when Met signaling is required, we turned to an ATP-competitive inhibitor of Met autophosphorylation, BMS777607 (Schroeder et al., 2009). We administered daily doses of BMS777607 to pregnant females to temporally inhibit Met signaling. In vehicle treated controls harvested at E17.5, the diaphragm is completely muscularized (n = 12/12; Fig. 5A). When BMS777607 was administered daily between E7.5 and E12.5, all embryos (n = 7/7) displayed bilateral dorsal muscleless patches and a ventral muscleless region (Fig. 5B). When treated at E8.5-E9.5 or E9.5-10.5 (Fig. 5C-D), all diaphragms had ventral muscleless regions (n = 14/14) and 35% had dorsal muscleless regions (n = 5/14). When treated at E11.5 and E12.5, after diaphragm progenitors have fully delaminated from the somites (Sefton et al., 2018), embryos had ventral muscleless regions (Fig. 5E-F; n = 8/8) and dorsal muscleless regions (n = 3/8). Quantification of the size of the ventral muscleless region indicates that all Met inhibition strategies lead to muscleless regions, with the largest muscleless region when Met is inhibited E7.5-E12.5 or E8.5-E9.5 (Fig. 5F). These data demonstrate that Met is continuously required from E7.5 to E12.5 for complete muscularization of the diaphragm. The regions requiring continuous Met signaling are on the leading edges of the diaphragm: the bilateral dorsal muscle and ventral midline muscle. These are the last regions to receive muscle progenitors that differentiate into myofibers.

**Figure 5.**
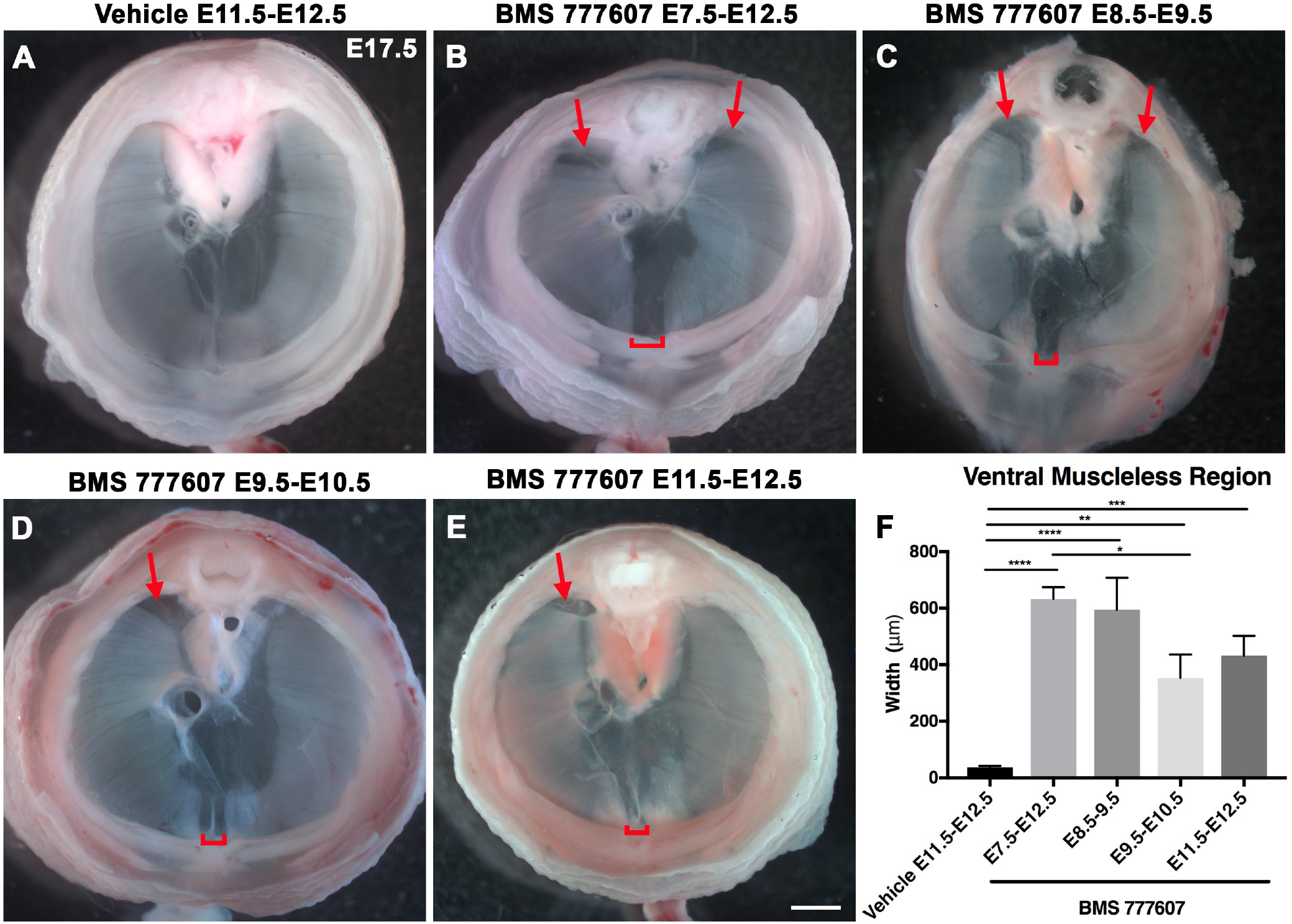
Reduction of Met signaling through inhibitor BMS777607 results in muscleless dorsal and ventral regions of the diaphragm. Unstained E17.5 diaphragms in cranial view. A) The left and right portions of the costal diaphragm meet in the ventral midline by E17.5 in vehicle treated controls. B-E) Dorsal left, dorsal right (arrows) and ventral midline regions (brackets) are muscleless when treated with BMS 777607 daily between E7.5-E12.5. H) Width of ventral midline muscleless region is significantly larger than vehicle treated controls when BMS777607 is administered at either early (E7.5-E8.5) or at later stages of diaphragm development (E11.5-E12.5). Significance tested with one-way ANOVA. * p<0.05, ** p<0.01, *** p<0.001, ****p<0.0001. Error bars represent standard error of the mean (SEM). Scale bar A-E: 1mm.

### Met signaling is required for survival, motility, and commitment of diaphragm muscle progenitors *in vitro*

While the regions most sensitive to Met inhibition are those that differentiate latest, it is unclear whether Met is required for proliferation, survival, differentiation, and/or motility. To investigate the function of Met in diaphragm muscle progenitors, we turned to an *in vitro* system to co-culture E12.5 diaphragm fibroblasts and myoblasts (Bogenschutz et al., 2020) in combination with BMS777607. PPFs were dissected from E12.5 *Pax3*^*CreKI/+*^; *Rosa*^*nTnG/+*^ embryos (Engleka et al., 2005; Prigge et al., 2013), in which *Pax3*-derived myogenic nuclei are GFP+ and PPF fibroblast nuclei are Tomato+, and cultured with either 10 μM BMS777607 or DMSO vehicle control. Overall, growth of GFP+ muscle progenitors was impaired with inhibitor treatment (Fig. 6A-C). To assess effects of the inhibitor on myoblast commitment, we examined *MyoD*. After 48 hours in culture, *MyoD* expression was reduced with inhibitor treatment (Fig. 6D). The percentage of cells co-expressing GFP and MyoD protein was similarly abrogated (Fig. 6E-F). Thus, Met inhibition not only reduces growth of myogenic cells, but it inhibits their commitment. By contrast, expression of the fibroblast marker *Gata4* was not significantly changed following inhibitor treatment (Fig. 1G). We tested whether the decreased growth of myogenic cells was due to decreased proliferation or increased apoptosis. Analysis of GFP+ myogenic cells labeled via EdU labeling indicates that BMS777607 treatment does not significantly change the percentage of proliferating cells (Fig. 6H-I). However, examination of apoptotic cells via staining for cleaved Caspase-3 showed that BMS777607 treatment significantly increased the number of apoptotic GFP+ myogenic cells (Fig. 6J-K). Met is also known to impair cell motility (reviewed by Birchmeier et al., 2003), and we found that the motility of GFP+ cells treated with BMS777607 was impaired, with reduced velocity and lower overall displacement (Fig. 6L-M). We also assessed effects on cell morphology by examining diaphragm muscle progenitors, labeled with membrane-bound GFP (via *Pax3*^*CreKI/+*^; *Rosa*^*mTmG/+*^). Cells were significantly more circular with BMS777607 treatment, which is consistent with compromised survival and motility (Fig. 6N-O). Overall, these data show that Met signaling is important for survival, motility, and commitment of diaphragm muscle progenitors.

**Figure 6.**
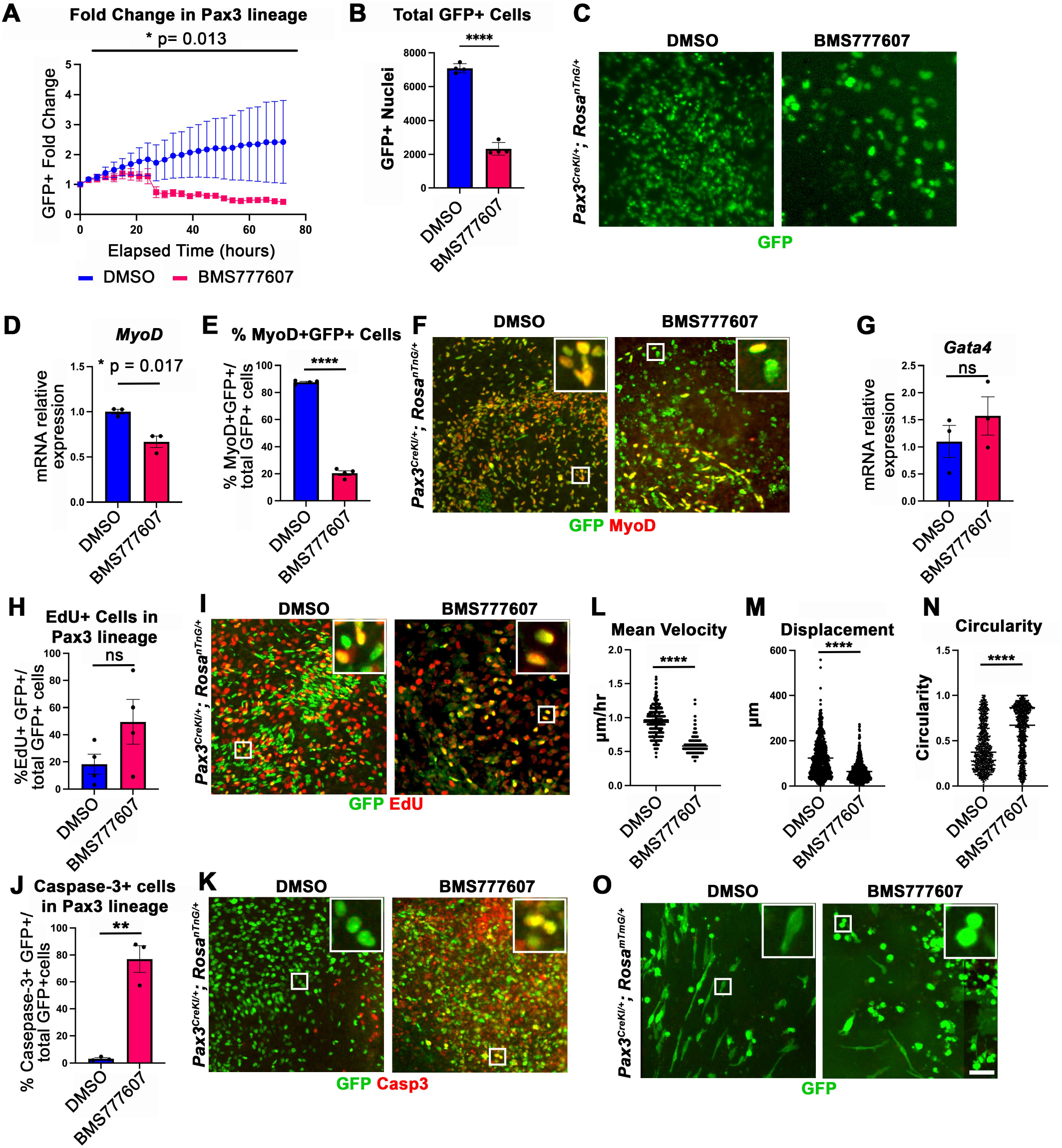
*In vitro* growth, motility, and differentiation of myogenic cells is impaired by pharmacological inhibition of Met signaling. A-0) E12.5 PPFs isolated from *Pax3*^*CreKI/+*^; *Rosa*^*nTnG/+*^ (A-M) or *Pax3*^*CreKI/+*^; *Rosa*^*mTmG/+*^ (N-O) embryos cultured with DMSO vehicle control or BMS 777607 and imaged on the ImageXpress Pico. A-C) The average ratio of GFP+ fold change (cell number at time T/cell number at time 0) ± SEM is plotted (A). The fold change (A) and total final count (B) of GFP+ cells are reduced following treatment with BMS777607 after 72 hr in culture (n = 4 biological replicates per treatment). Representative GFP images shown in C. D-F) *MyoD* expression (via qPCR, D) and percentage of MyoD+GFP+ cells (E) is significantly lower after 48 h in culture with Met inhibition. Representative images of GFP and MyoD expression (F). G) Expression of PPF fibroblast marker *Gata4* is not significantly affected by BMS777607 treatment. H-I) EdU-labeling of GFP+ cells is not significantly changed by BMS777607 treatment. J-K) Cleaved Caspase-3 expression was significantly increased with BMS777607 treatment. L-M) Mean velocity and total displacement of GFP+ nuclei were decreased with BMS777607 treatment. GFP+ nuclei were imaged every 8 minutes over 14 h to track cell motility. N-O). BMS777607-treated GFP+ cells were significantly more circular. Representative images (O). * p<0.05, ** p<0.01, **** p<0.0001. Statistical changes in cell number over time were determined using repeated measure ANOVA on the log2 transformed fold change over time. Error bars represent standard error of the mean (SEM). Scale bar: 100 μm.

### Met signaling is required for population of muscle progenitors at the diaphragm’s leading edges and the consequent development of the dorsal and ventral-most muscle regions

Our *in vivo* studies show that HGF/Met signaling is required for development of the dorsal and ventral-most regions of the diaphragm muscle and our *in vitro* studies find that Met is required for muscle progenitor survival, motility, and commitment. Based on these data, we hypothesized that *in vivo* loss of dorsal and ventral muscle regions was due to fewer muscle progenitors and/or myoblasts at the dorsal and ventral leading edges of the diaphragm when it is expanding. To test this, wild-type embryos were treated with BMS777607 daily E7.5-E11.5, harvested at E12.5, and stained for myogenic cells with a cocktail of Pax7, MyoD, and Myosin antibodies as well as for EdU, cleaved Caspase-3, and Neurofilament. We found that the PPFs (identified and outlined in 3 dimensions by their unique morphology, viewed by autofluorescence) were more variable in size, but not significantly decreased in size from control diaphragms. We also found, consistent with our analysis of *HGF*^*Δ/Δ*^ and *Met*^*Δ/Δ*^ mice, that nerve branching was strongly reduced by the inhibitor (Fig. 7 C, F, J, M). Supporting our hypothesis, the inhibitor led to a reduction in the number of mononuclear progenitors and myoblasts at the ventral and dorsal leading edges of the muscle (arrows in Fig. 7A, D, H, K). Inhibitor treated embryos also showed reduced numbers of EdU+ cells overall (Fig. 7B, E, G) and an increased number of cleaved Caspase-3 positive cells within the PPFs (Fig. 7I, L, N). Thus these data demonstrate that *in vivo* Met signaling is required to promote proliferation and survival of myogenic cells, and its inhibition leads to a loss of muscle progenitors and myoblasts at the leading edges and results in a loss of the dorsal-most and ventral-most diaphragm muscle.

**Figure 7.**
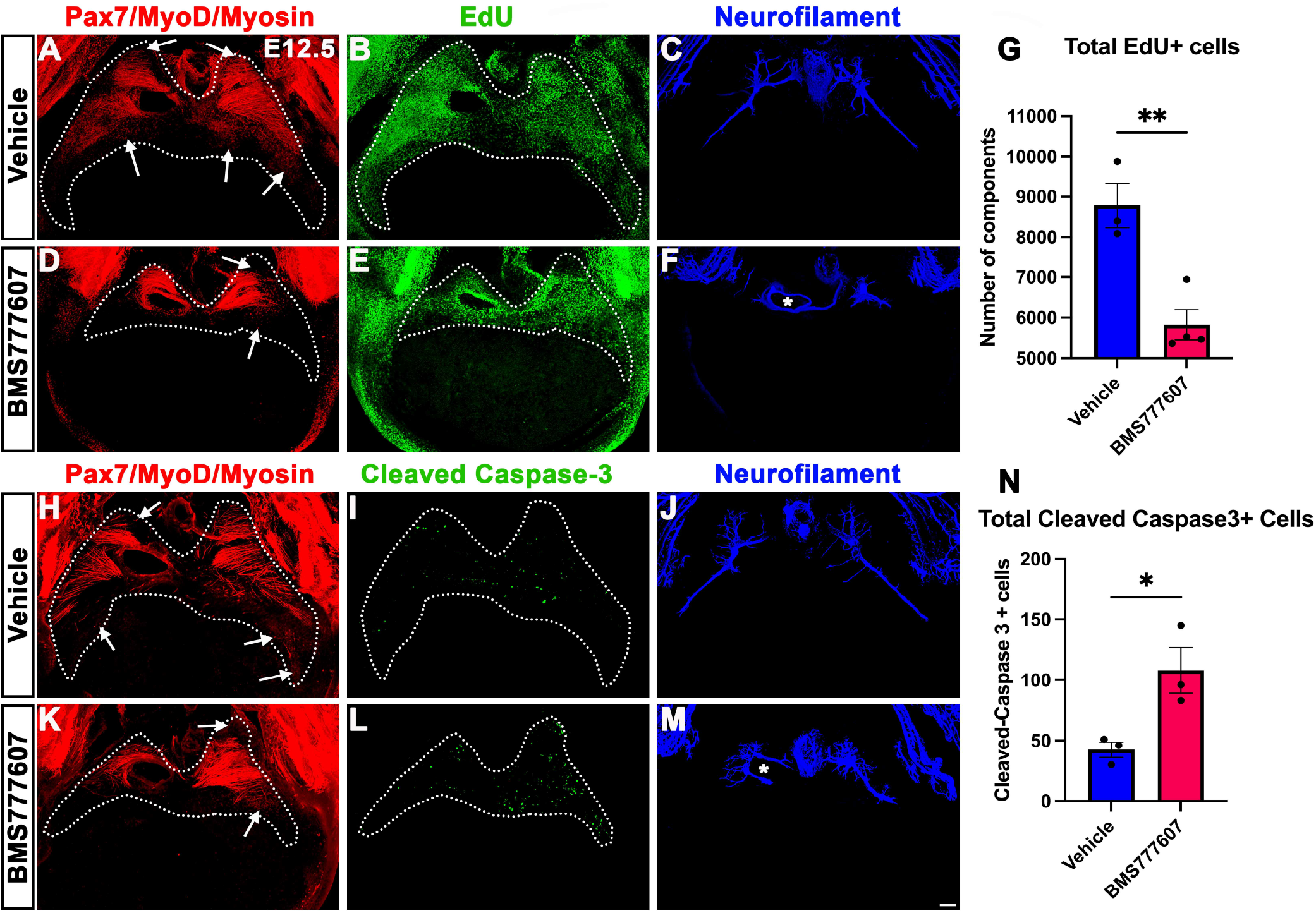
Inhibition of Met signaling *in vivo* alters cell proliferation, apoptosis and phrenic nerve morphology. Embryos were treated with BMS777607 or vehicle control daily between E7.5-E11.5, harvested at E12.5, stained for Pax7, MyoD, and Myosin, and imaged in whole mount cranial view on the confocal. Treatment with BMS777607 leads to fewer mononuclear myogenic cells on the dorsal and ventral leading edges of the diaphragm (arrows; A, D, H, K), reduced total EdU positive nuclei (B, E, G), increased cleaved-caspase-3 positive (I, L, N), and aberrant phrenic nerve branching (F, M), where the right phrenic neve wraps around vena cava (asterixis in F, M). Scale bar: 100 μm.

## Discussion

The diaphragm is an essential mammalian skeletal muscle, playing a critical role in respiration and serving as a barrier that separates the thorax from the abdomen. Not only is the diaphragm a functionally important muscle, but it serves as an excellent system to study muscle patterning and morphogenesis, since it is a flat muscle that largely develops in two dimensions. Development of the diaphragm, like other skeletal muscles, requires the integration of muscle, connective tissue, and nerve that arise from different embryonic sources. Our study establishes that the connective tissue fibroblasts are the source of a molecular signal, HGF, that coordinates development of muscle and nerve (Fig. 8).

**Figure 8.**
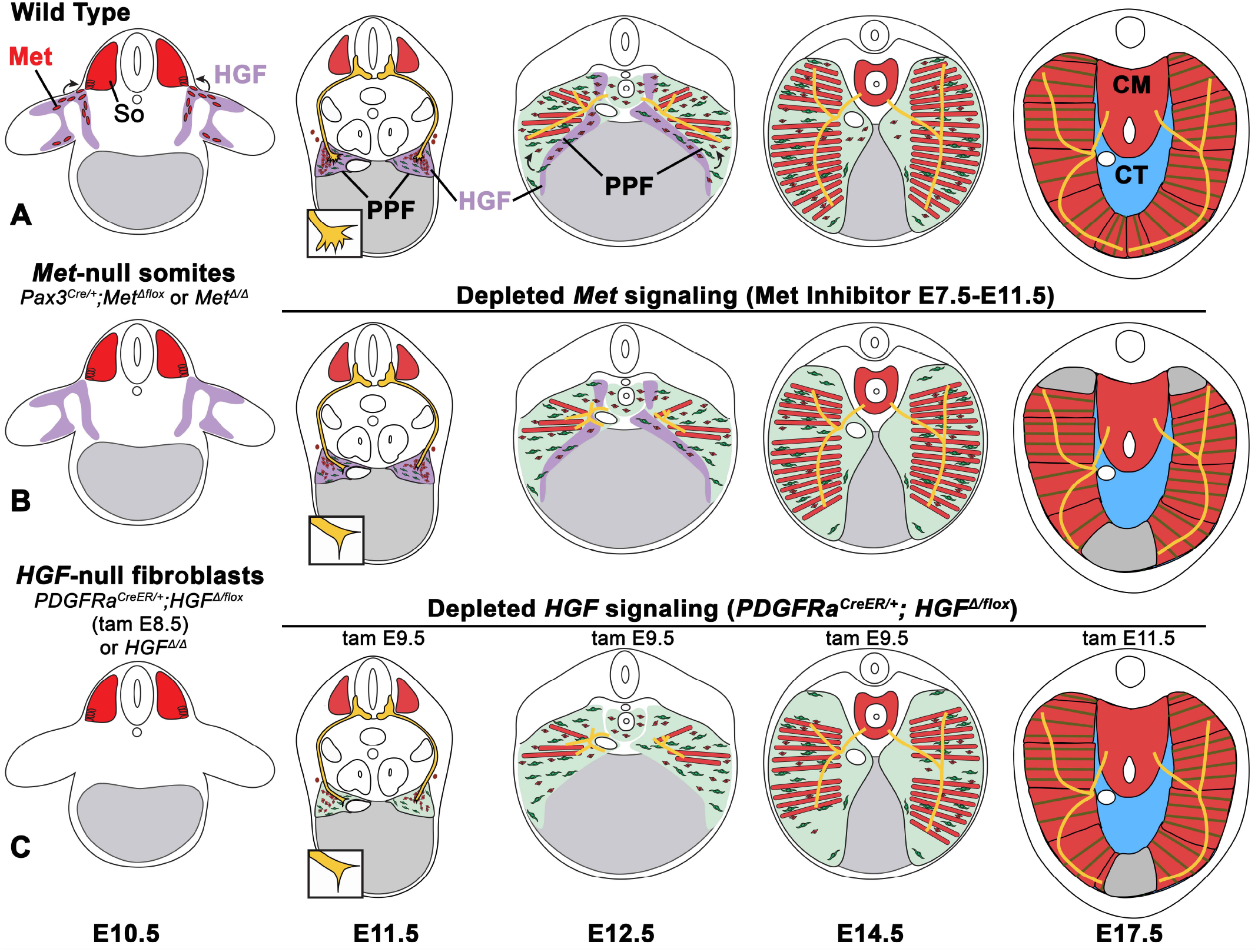
Model of HGF-MET signaling in skeletal muscle and phrenic nerve in the diaphragm. Our data support a model where fibroblast-derived HGF and muscle progenitor-derived MET are critical for muscularization of the limb and diaphragm. Null mutations or tissue-specific early mutations (prior to E9.0) lead to muscleless limb and diaphragm and reduced defasciculation of the phrenic nerve. Later mutations or incomplete depletion lead to partial muscularization, where the nerve displays aberrant branching that tracks with muscle localization. B) Met depletion through chemical inhibition between E7.5-E11.5 results in reduced muscle progenitors at the leading edge of the diaphragm at E12.5 and muscleless regions in the dorsal and ventral diaphragm at E17.5. *In vitro* and *in vivo* data suggest these effects are mediated by increased apoptosis and reduced motility. C) Early mutations in fibroblast-derived HGF (tamoxifen administration at E9.5) result in large muscleless regions at E14.5, while later mutations (tamoxifen administration at E11.5) affect the leading ventral edge of the diaphragm. PPFs, pleuroperitoneal folds; So, somite; NT, neural tube.

Development of muscle and its innervating motor neurons must be tightly integrated to produce a functional muscle. The connective tissue is an ideal candidate tissue to orchestrate this process as it enwraps myofibers and neuromuscular junctions (Nassari et al., 2017; Sefton and Kardon, 2019). In the diaphragm, the PPFs are the source of the diaphragm’s connective tissue fibroblasts and critical for overall diaphragm morphogenesis (Merrell et al., 2015). Thus, the PPFs are likely to coordinately regulate muscle and the phrenic nerve. A number of previous studies also suggested that HGF/MET is a key signaling pathway for this coordination since it has been found to regulate both muscle development and innervation (Bladt et al., 1995; Dietrich et al., 1999; Ebens et al., 1996; Lamballe et al., 2011; Maina et al., 1996; Yamamoto et al., 1997). In addition, previous studies established that *Met* is required for diaphragm development (Bladt et al., 1995; Dietrich et al., 1999; Maina et al., 1996). Here we used conditional mutagenesis to specifically target *HGF* and *Met* in PPF-derived connective tissue fibroblasts or muscle progenitors to genetically dissect the role of these cells and HGF/MET signaling in diaphragm development (Fig. 8). First, as expected we found that HGF/MET signaling is required for the initial delamination and migration of Met+ muscle progenitors from the somites to the nascent diaphragm. More surprisingly, we found that HGF expressed at the dorsal and ventral margins of the expanding PPFs is continuously required for proliferation, survival, and motility of muscle progenitors and the consequent expansion and full muscularization of the diaphragm. Our experiments also showed, that unlike the muscle progenitors, HGF/MET signaling is not required for the recruitment and targeting of phrenic nerve axons to the nascent diaphragm. However, PPF-derived HGF is crucial for the defasciculation and primary branching of the nerve, independent of muscle. Subsequently, PPF-derived HGF is required, likely muscle-dependent, for outgrowth of the primary branches of the phrenic nerve axons throughout the diaphragm muscle. Previous studies have identified how phrenic motor neurons are specified in the motor column (Dasen et al., 2008; Dasen et al., 2003; Dasen et al., 2005; Jung et al., 2010; Rousso et al., 2008) and later arborize (Philippidou et al., 2012) to form neuromuscular junctions (Burden, 2011; Li et al., 2008; Wang et al., 2003; Yumoto et al., 2012). Only one study (Uetani et al., 2006) has identified molecular regulators, Receptor Protein Tyrosine Phosphatases and, of phrenic nerve defasciculation and primary axon outgrowth, and our study now adds HGF/MET as another regulator of this process. Altogether our study demonstrates that PPF-derived connective tissue fibroblasts and HGF control several important aspects of diaphragm muscle and phrenic nerve development. Connective tissue and HGF are also likely to coordinately regulate the development of other muscles and their motor neurons, as both connective tissue and HGF/MET signaling have been implicated in the development and innervation of limb and back muscles (Caruso et al., 2014; Helmbacher, 2018).

Our study also provides insights into how the diaphragm might have evolved. The diaphragm muscle is unique to mammals and how it evolved in mammals is a major unanswered question (Perry et al., 2010). Evolution of the diaphragm involved the acquisition of developmental innovations in mammals that are absent from birds and reptiles. Comparison of these groups suggests the following important developmental innovations: formation and expansion of PPFs across the liver to separate the thoracic and abdominal cavities, recruitment of muscle progenitors to the PPFs and their expansion and differentiation into the radial array of myofibers, and recruitment and targeting of motor neurons to the diaphragm muscle (Hirasawa et al., 2015; Hirasawa and Kuratani, 2013; Sefton et al., 2018). HGF/MET signaling has important, conserved functions in the development of most vertebrate hypaxial muscles (Adachi et al., 2018; Haines et al., 2004; Okamoto et al., 2019). Here we identify that *HGF* expressed by the PPFs is crucial for recruitment and expansion of diaphragm muscle progenitors. Our experiments conditionally deleting *HGF* also indicate that migration of diaphragm and shoulder (spinodeltoid and acromiodeltoid) muscle progenitors are *HGF*-dependent and occur contemporaneously. Recruitment of shoulder muscle progenitors to the nascent diaphragm has been proposed as important for the evolutionary origin of the mammalian diaphragm (Hirasawa and Kuratani, 2013). Thus our experiments suggests that the evolutionary acquisition of *HGF* expression in the PPFs may have been a key event that allowed a subset of shoulder muscle progenitors to be recruited to the nascent diaphragm. Our experiments also indicate that continued *HGF* expression is critical for the full muscularization of the diaphragm. Therefore, once *HGF* was expressed in the PPFs there may have been selection for its continued expression to enable expansion of the muscle and complete separation of the thoracic and abdominal cavities. Interestingly, our data indicate that the evolutionary acquisition of *HGF* expression in the PPFs is not sufficient to recruit motor neurons and so some other signal(s) must be involved in the evolutionary recruitment of the phrenic nerves. Also still unknown are the developmental and evolutionary mechanisms driving formation and morphogenetic expansion of the PPFs.

Finally, our study elucidates some of the cellular mechanisms underlying the etiology of CDH. CDH is characterized by defects in the muscularization of the diaphragm, and two sites where muscle is commonly absent are the dorsal-most region of the diaphragm (designated posterior in humans and hernias in this area are called Bochdalek hernias) and the ventral-most diaphragm (anterior in humans and hernias here are called Morgagni hernias) (Ackerman et al., 2012; Irish et al., 1996; Kardon et al., 2017). Our analysis of diaphragms in which HGF/MET signaling is perturbed has found that these two regions are the most likely to have muscularization defects and suggests mechanistically why the dorsal-most and ventral-most diaphragm are most susceptible to muscularization defects. Mutations or variants in any gene or signaling pathway that, similar to HGF/MET, regulates the proliferation, survival, or motility of muscle progenitors will lead to a depletion of the pool of muscle progenitors and the consequent loss of the dorsal and ventral-most diaphragm muscle, since these regions develop last (Fig. 8). Most surprisingly, our analysis revealed that the muscleless connective tissue regions in mice with deletion of *HGF* in the PPFs or pharmacological inhibition of MET signaling do not herniate. We previously conducted a detailed analysis of mice in which the transcription factor *Gata4* was deleted in the PPFs (Merrell et al., 2015). In these mice muscleless connective tissue regions develop, but these regions always herniate and give rise to herniated tissue covered by a connective tissue sac. Based on our study of *Gata4*, we had proposed that such “sac” hernias (Pober, 2007) develop when localized regions of amuscular tissue develop in juxtaposition with muscularized tissue; the biomechanical difference in strength between these regions allows the abdominal tissues to herniate through the weaker amuscular regions. However, the lack of herniation in the *HGF* mutants demonstrates that the formation amuscular connective tissue regions is not sufficient to cause “sac” hernias. While formation of amuscular regions is likely a critical step in the formation of “sac” hernias, other defects in connective tissue integrity are likely necessary to cause these susceptible amuscular regions to actually herniate. Thus, herniation may be a multi-step process involving loss of muscle followed by defects in connective tissue strength or elasticity that allow the liver to herniate into the thoracic cavity. Further insight into these processes will be critical for the development of therapeutic targets in the future.

## Materials and Methods

### Mice and staging

All mouse lines have been previously published. We used *Prx1Cre*^*Tg*^ (Logan et al., 2002), *PDGFRa*^*CreERT2*^ (Chung et al., 2018), *Pax3*^*Cre*^ (Engleka et al., 2005), *Pax7*^*iCre*^ (Keller et al., 2004), *Pax7*^*CreER*^ *(Murphy et al., 2011)* and *Hprt-cre (Tang et al., 2002)* Cre alleles. Cre-responsive reporter alleles included *Rosa26*^*LacZ*^ (Soriano, 1999), *Rosa*^*nTnG*^ (Prigge et al., 2013), and *Rosa*^*mTmG*^ (Muzumdar et al., 2007). We used *HGF*^*fl*^ (Phaneuf et al., 2004) and *Met*^*fl*^ (Huh et al., 2004) conditional alleles. *HGFα*^*Δ/+*^ and *Met*^*Δ/+*^ mice were generated by breeding *HGF*^*fl*^ and *Met*^*fl*^ mice to *Hprt-cre* mice. Embryos were staged as E0.5 on the day dams presented with a vaginal plug. Mice were backcrossed onto a C57/Bl6J background. Experiments were performed in accordance with protocols approved by the Institutional Animal Care and Use Committee at the University of Utah.

### Immunohistochemistry, immunofluorescence, and in situ hybridization

For section immunofluorescence, OCT (optimal cutting temperature)-embedded tissues were tissues were sectioned to 10μm thickness and fixed for 5 minutes in 4% paraformaldehyde (PFA). Tissue sections were blocked for 60 minutes in 5% goat serum in PBS, incubated overnight at 4°C in primary antibodies. Sections were washed in PBS, incubated with secondary fluorescent antibodies (used at 1-5 μg/ml; Jackson Laboratories or Thermo Fisher) for 2 h at RT, washed with PBS, stained for 5 minutes with Hoechst to label nuclei, post-fixed in 4% PFA, rinsed in water and mounted with Fluoromount-G (Southern Biotech). Primary antibodies are listed in supplemental Table 1. Sections were imaged on an Olympus BX63.

For immunofluorescence on PPF cell cultures, cells were fixed in 4% PFA for 20 minutes at room temperature, washed in PBS, blocked for 60 minutes in 5% goat serum with 0.1% Triton X-100 in PBS, and stained overnight for primary antibodies (see Supplementary Table 1). Cells were then washed in PBS, incubated for 2 hours in secondary antibodies, washed in PBS, incubated in Hoechst to label nuclei, washed in PBS, rinsed in water and mounted in Fluoromount. EdU (Life Technologies) was applied to cells 1 hour prior to fixation and detected after secondary labeling based on the manufacturer’s instructions with Alexa647 picolyl azide. Stained cells were imaged with ImageXPress Pico automated cell imager (Molecular Devices).

Wholemount embryos were fixed for 24 h in 4% PFA at 4°C, dissected, incubated for either 2hrs at RT or overnight at 4°C in Dent’s bleach (1:2 30% H_2_O_2_:Dent’s fix) and stored in Dent’s fix (1:4 DMSO:methanol) for at least 5 days at 4°C. Embryos were washed in PBS, blocked for 1 h in 5% goat serum and 20% DMSO, incubated in primary antibodies (see Supplementary Table 1) for 48 h, washed in PBS, incubated in secondaries for 24-48 h, washed in PBS and cleared BA:BB (33% benzyl alcohol, 66% benzyl benzoate) at RT. Embryos labeled with AP-conjugated anti-myosin were heat-inactivated at 65°C for 1 h, incubated in primary antibody for 48 h, and detected with 250 μg/ml NBT and 125 μg/ml BCIP (Sigma) in alkaline phosphatase buffer. For detection of HRP-conjugated secondary antibodies, embryos were incubated in 10 mg Diaminobenzidine tetrahydrochloride in 50mL PBS with 7 μl hydrogen peroxide for approximately 20 min.

For wholemount EdU analysis in embryos, 10 μg/g of body weight of EdU was administered to pregnant females 1 h prior to harvest via IP injection.

Wholemount in situ hybridization was performed as previously described (Riddle et al., 1993). For wholemount β-galactosidase staining, embryos were fixed overnight in 1% PFA at 4°C and 2 mM MgCl_2_. Diaphragms were dissected, washed in PBS and then in LacZ rinse buffer (100mM sodium phosphate, 2mM MgCl_2_, 0.01% sodium deoxycholate and 0.02% Igepal) and stained for 16 h at 37°C in X-gal staining solution (5mM potassium ferricyanide, 5 mM potassium ferrocyanide and 1mg/ml X-gal).

### Microscopy and three-dimensional rendering

Wholemount fluorescent images were taken on a Leica SP8 confocal microscope. Optical stacks of wholemount images were rendered and structures highlighted using Fluorender (Wan et al., 2009). To highlight features (such as the phrenic nerve in Fig. 4), objects were selected in Fluorender based on morphology on individual Z optical sections using the paint brush tool, and then these objects were extracted, rendered, and pseudo-colored. For EdU and cleaved Caspase-3 labeling, PPFs were first selected based on morphology and the total PPF area measured. Individual nuclei from PPFs were then counted using the Component Analyzer Tool.

### Cell culture, media and reagents

E12.5 embryos were dissected from pregnant *Rosa*^*mTmG/mTmG*^ or *Rosa*^*nTnG/nTnG*^ females mated with *Pax3*^*CreKI/+*^ males. Embryos and PPFs were dissected as previously described (Bogenschutz et al., 2020). Growth of both PPF fibroblasts and myogenic cells was promoted using DMEM/F-12 GlutaMAX (Invitrogen), 10% FBS, 50 μg/mL gentamycin, and 0.5nM FGF. PPFs were grown in a 37°C incubator overnight and then imaged on an ImageXPress Pico automated cell imager (Molecular Devices), for 1-4 d, changing media in the wells every 2 d.

### Chemical treatments

For BMS777607 administration to pregnant females, 0.05mg/g of body weight (e.g., 1mg BMS777607 for a 20g mouse) in 70% PEG-300 was administered via oral gavage. Vehicle alone (70% PEG300 in PBS) was administered to control pregnant dams. For cell culture experiments, BMS777607 was used at 10 μM concentration in 0.001% DMSO, and vehicle controls were treated with 0.001% DMSO.

### Cell growth, motility and shape analysis

Proliferating myogenic cells were imaged using the ImageXPress Pico which took GFP, Tomato and phase images every 3 h of the entire PPF sample for 72 h total.

CellReporterXpress (Molecular Devices) software was then used to count GFP+ cells per time point to calculate growth of myogenic cells over time. To control for differences in initial number of myogenic cells per well, fold changes of cell growth were calculated by dividing each treatment by the initial cell number at time 0. For cell motility analysis of *Pax3*^*CreKI/+*^; *Rosa*^*nTnG/+*^ nuclei, GFP and phase images were taken every 8 minutes for 14 h. Tracking, cell velocity and displacement of peripheral cells (as an analogue for the leading edge of the PPFs) were determined using TrackMate (Tinevez et al., 2017). For cell shape analysis on *Pax3*^*CreKI/+*^; *Rosa*^*mTmG/+*^ membranes, GFP and phase images were imaged every 4 minutes apart for 68 minutes total. Circularity of peripheral cells was analyzed in Fiji.

### RNA extraction, cDNA synthesis, and quantitative polymerase chain reaction (qPCR)

The *Quick*-RNA Microprep Kit (Zymo, Irvine, CA) was used to extract total RNA according to manufacturer’s protocol. Applied Biosystems High-Capacity RNA-to-cDNA kit (Thermo-Fisher) was used to synthesize cDNA from purified RNA according to manufacturer’s protocol. qRT-PCR was used to analyze expression of *Met, Pax7, HGF, MyoD1* and *Gata4* using pre-validated primer sets (TaqMan, ThermoFisher; Supplemental Table 2). 10 μl reaction volumes were prepared using TaqMan Fast Advanced Master Mix (ThermoFisher). The following conditions were used for amplification: 20 s at 95°C followed by 40 cycles at 95°C for 1 s and 60°C for 20 s. Gene expression levels were normalized against *18S* ribosomal RNA for each sample and fold changes calculated using 2^-*Δ/ΔCt*^ method (Schmittgen and Livak, 2008) by setting expression levels of each gene in DMSO-treated cell culture as 1. Data from 3 biological replicates were calculated and plotted as average fold changes with standard error of the mean (SEM).

### Statistical analysis

Data are presented as ±SEM. For growth comparison between chemical treatments, repeated ANOVA analysis was run on the log2 fold change of GFP+ cells to normalize the distribution of cell growth over time. Unpaired two-tailed t-tests were used for other statistical analyses.

## Acknowledgements

We thank B. Collins for critical reading of the manuscript and Y. Wan for help with Fluorender analysis.

## Competing interests

No competing interests declared.

## Funding

This work was supported by the National Institutes of Health [R01HD087360 to G.K., F32 HD093425 and 1K99HD101682 to E.M.S.] and Wheeler Foundation to G.K.

## Figure Legends

**Figure S1.**
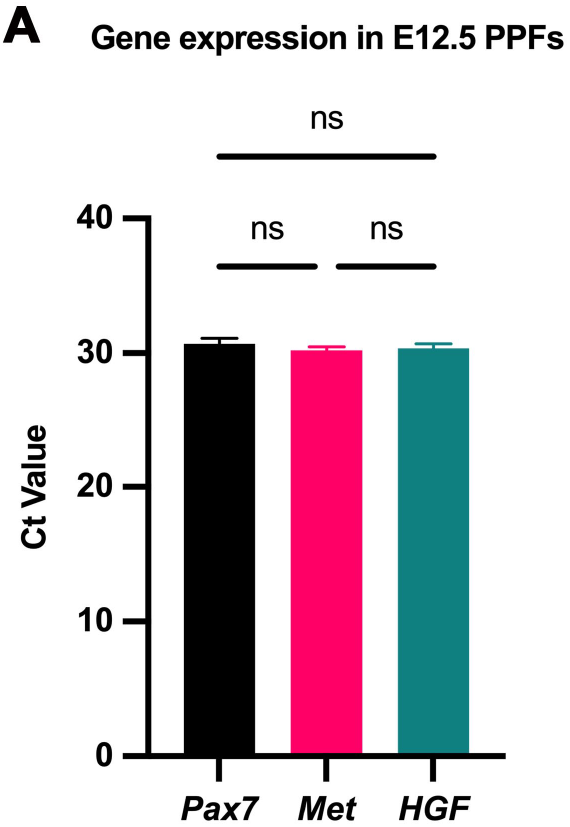
*Met* is expressed in the diaphragm at E12.5. Gene expression assayed by qPCR on PPFs isolated from E12.5 WT+/+ embryos. Cycle threshold values of *Pax7, Met*, and *HGF* are comparable at this stage (no significant differences, as tested by one-way ANOVA).

**Figure S2.**
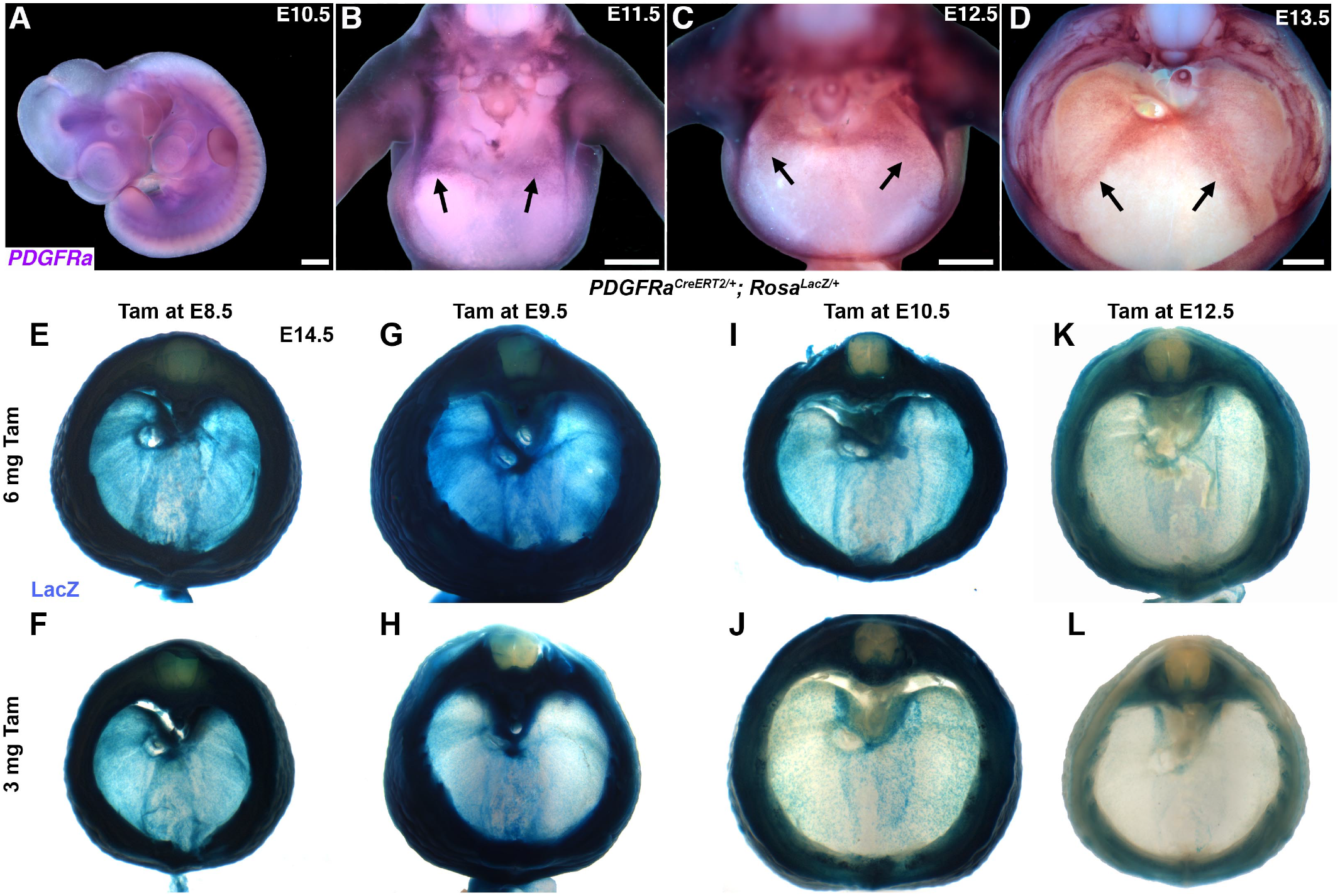
*PDGFRα* is expressed within the developing diaphragm and *PDGFR*^*CreERT2*^ allele targets the PPFs. (A-D) mRNA *in situ* hybridization for *PDGFRα*at E10.5 in lateral view (A); E11.5 (B), E12.5 (C) and E13.5 (D) are in cranial view with dorsal to the top, dissected to diaphragm with the heart and lungs removed. Arrows indicate the leading edges of the PPFs. Scale bars: 500 μm. (E-L) *PDGFRα*^*CreERT2/+*^ labels PPF fibroblasts, including muscle connective tissue and central tendon, but not muscle. As expected, fibroblasts throughout the body wall are also labeled. Approximately E14.5 *PDGFRα*^*CreERT2/+;*^ *Rosa*^*LacZ/+*^ embryos stained for LacZ, given different doses tamoxifen at either E8.5, E9.5, E10.5 or E12.5 (right column). Fibroblast labeling is stronger at higher doses of tamoxifen (6mg) and fibroblasts are not strongly labeled with tamoxifen at E12.5.

**Figure S3.**
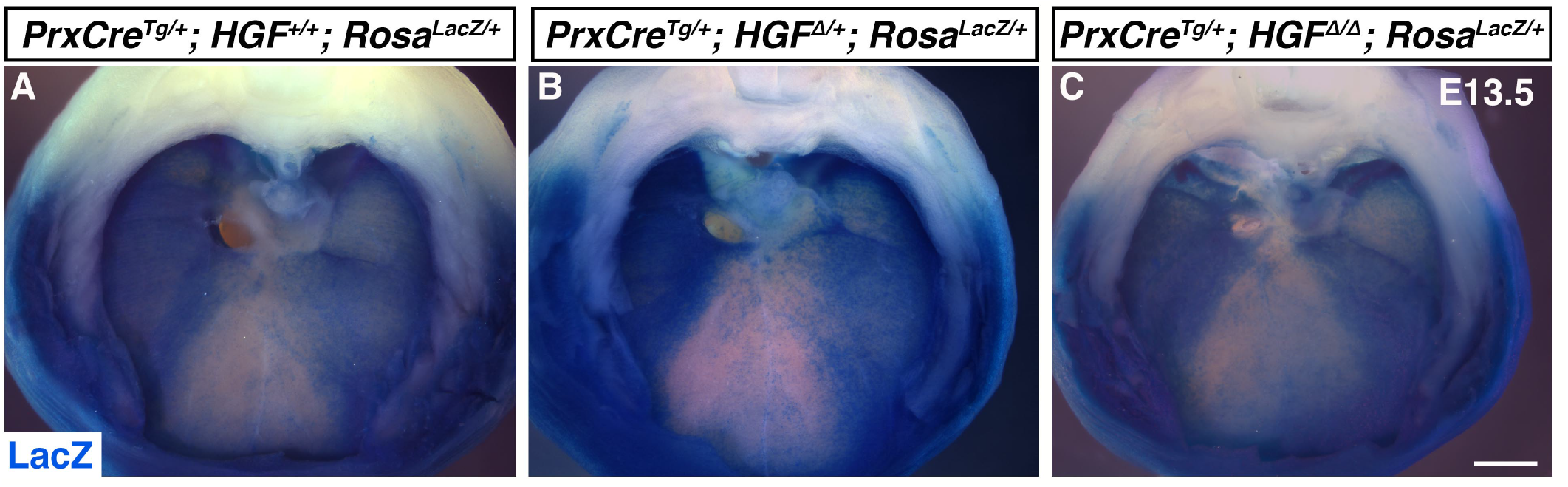
Pleuroperitoneal folds continue to spread normally following deletion of *HGF*. A-C) Cranial view of E13.5 *Prx1Cre*^*Tg/+*^; *Rosa*^*LacZ/+*^ diaphragms stained for LacZ. The *Prx1* transgene labels the PPFs (Merrell et al., 2015). A) By E13.5, the PPFs have spread throughout the dorsal and all but the ventral most region of the diaphragm. B-C) Fold spreading by E13.5 is unaffected in embryos heterozygous or null for *HGF*. Scale bar: 500 μm.

**Figure S4.**
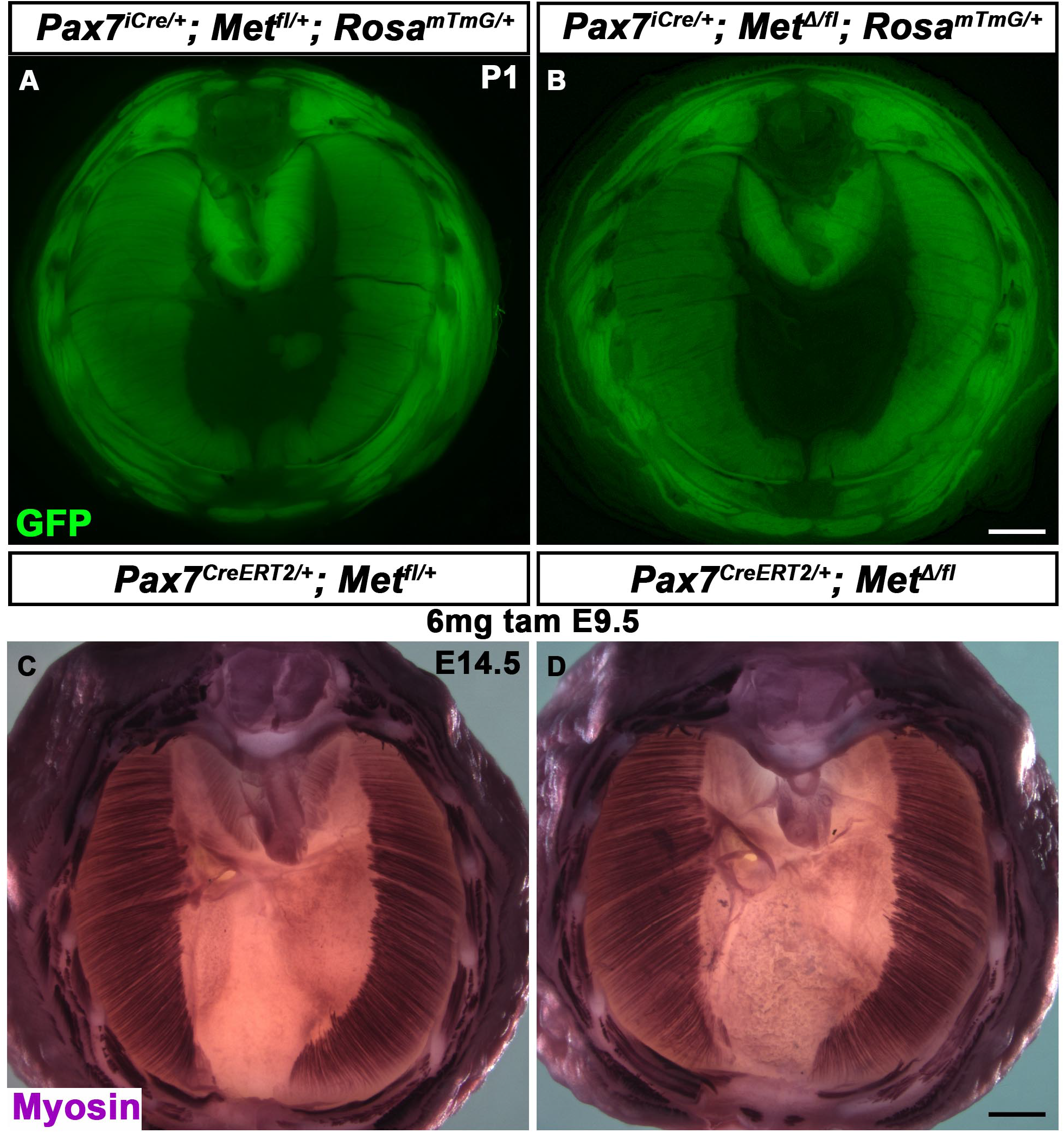
Normal muscularization of the diaphragm following deletion of *Met* in the *Pax7* lineage. (A-B). Image of endogenous GFP fluorescence in musculature of P1 diaphragms. (C-D) Myosin staining of diaphragm musculature at E14.5 with inducible *Pax7*^*CreERT2/+*^ given 6mg of tamoxifen at E9.5. Scale bar (A-B): 1 mm; (C-D): 500 μm.

**Supplemental Table 1**

**Antibodies used in study**

**Table S2. Oligonucleotides for qPCR (related to Figure 6)**.

## Notes

### Competing Interest Statement

The authors have declared no competing interest.

